# Interpretable machine learning of action potential duration restitution kinetics in single-cell models of atrial cardiomyocytes

**DOI:** 10.1101/2022.05.13.491795

**Authors:** Euijun Song, Young-Seon Lee

## Abstract

Action potential duration (APD) restitution curve and its maximal slope (Smax) reflect single cell-level dynamic instability for inducing chaotic heart rhythms. However, conventional parameter sensitivity analysis often fails to describe nonlinear relationships between ion channel parameters and electrophysiological phenotypes, such as Smax. We explored the parameter–phenotype mapping in a population of 5,000 single-cell atrial cell models through interpretable machine learning (ML) approaches. Parameter sensitivity analyses could explain the linear relationships between parameters and electrophysiological phenotypes, including APD_90_, resting membrane potential, Vmax, refractory period, and APD/calcium alternans threshold, but not for Smax. However, neural network models had better prediction performance for Smax. To interpret the ML model, we evaluated the parameter importance at the global and local levels by computing the permutation feature importance and the local interpretable model-agnostic explanations (LIME) values, respectively. Increases in I_CaL_, I_NCX_, and I_Kr_, and decreases in I_K1_, I_b,Cl_, I_Kur_, I_SERCA_, and I_to_ are correlated with higher Smax values. The LIME algorithm determined that INaK plays a significant role in determining Smax as well as Ito and I_Kur_. The atrial cardiomyocyte population was hierarchically clustered into three distinct groups based on the LIME values and the single-cell simulation confirmed that perturbations in I_NaK_ resulted in different behaviors of APD restitution curves in three clusters. Our combined top-down interpretable ML and bottom-up mechanistic simulation approaches uncovered the role of I_NaK_ in heterogeneous behaviors of Smax in the atrial cardiomyocyte population.

## 1. Introduction

Heart is believed as a complex dynamical system with the presence of nonlinearity and multistability, particularly in pathological conditions such as cardiac arrhythmias [1]. Spatiotemporal chaotic behaviors of the electrical wave propagation can lead to intractable or life-threatening cardiac arrhythmias such as atrial fibrillation (AF) or ventricular fibrillation (VF) [2]. This nonlinear nature of cardiac electrophysiology is originated from the interactions between subcellular molecular/ion dynamics and tissue-level electrical wave dynamics [3]. One important concept to explain single cell-level dynamic instability is an action potential duration (APD) restitution curve, which represents the adaptation properties of action potential to the heart rate. Additionally, the presence of rate-dependent alternans in APD or calcium transient (CaT) is related to cellular instability [4]. Both the maximal slope (Smax) of the APD restitution curve and the presence of APD/calcium alternans are known to be closely linked to the instability of spiral wave dynamics, eventually leading to AF or VF [5, 6]. Recently, many computational models have been developed to quantitatively analyze the nonlinear phenomenon of cardiac electrophysiology [7]. Additionally, patient-specific multiscale heart models are being used to understand the complex dynamics of cardiac arrhythmias and to personalize ablation/drug treatments [8–10]. These computational cardiac models, however, do not fully capture disease-specific or patient-specific electrical/structural remodeling states. Therefore, it is still challenging to determine quantitative relationships between individual ion channel profiles and dynamic electrophysiological phenotypes such as APD restitution kinetics due to the highly nonlinear and multiscale nature of the cardiac system.

Quantitative prediction of phenotypes from individual parameter profiles is critical to evaluating the complex disease risk and pharmacological responses at patient-specific and single-cell levels [11, 12]. While some phenotypes are robust to parameter perturbations, most complex traits usually show nonlinear behaviors depending on parameter combinations [13, 14]. Conventional parameter sensitivity analysis can describe linear relationships between parameters and quantitative phenotypes based on the linear approximation of phenotypes in the local neighborhood of a given baseline [15, 16]. However, linear models cannot fully explain the nonlinear behaviors of complex dynamical systems on global parameter space. One powerful tool to explore the nonlinear parameter–phenotype relationships on a global parameter space is machine learning (ML). ML can provide a highly efficient tool for predicting phenotypes from parameter combinations, bypassing the enormous computational cost of mechanistic computational simulations [17, 18]. However, ML models are commonly referred to as black boxes due to their ambiguous internal mechanisms of the prediction process. Recently, several model-agnostic interpretation methods have been developed to interpret various ML models [19]. Similar to the parameter sensitivity analysis, parameter importance can be evaluated from trained ML models by computing global or local feature importance measures, from which we can better understand parameter–phenotype relationships at both global and local levels on the parameter space [19, 20].

In this study, we explore the mapping between ion channel parameters and quantitative electrophysiological phenotypes in single-cell computational models of human atrial cardiomyocytes using parameter sensitivity analysis and ML. We generate a large parameter– phenotype mapping dataset from *bottom-up* mechanism-based atrial cell models by extensively perturbing ion channel parameters. Then, we train various ML models to capture the nonlinear parameter–phenotype relationships, focusing on the Smax. To assess how parameters contribute to phenotypes, we calculated global/local parameter importance measures using ML interpretation methods. Finally, we perform *in-silico* perturbation experiments in atrial cell models to confirm the findings from the *top-down* ML analyses.

## 2. Methods

### 2.1. Computational modeling of atrial cardiomyocytes

We generated a population of single-cell computational models using the Grandi human atrial cell model [21] by perturbing 18 key model parameters including 14 transmembrane ionic currents (I_Na_, I_b,Na_, I_CaL_, I_b,Ca_, I_to_, I_K1_, I_Kr_, I_Ks_, I_Kur_, I_Kp_, I_ClCa_, I_b,Cl_, I_NaK_, I_NCX_) and 4 calcium handling parameters (I_SR,release_, I_SR,leak_, I_SERCA_, I_SLCaP_). Briefly, the cell model was described as the following differential equations:

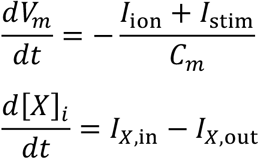

where V_m_ is the transmembrane potential, I_ion_ is the total ionic current, I_stim_ is the stimulus current, C_m_ is the total membrane capacitance, [*X*]_*i*_ is the intracellular concentration of component X (e.g., Na^+^, K^+^, Ca^2+^), and I_x,in(or out)_ is the inward (or outward) current related to X. The ion currents are typically nonlinear functions of the transmembrane potential, intracellular ion concentration, and gating state variables, which were constructed from the experimental voltage-clamp data. The biophysical details can be found in Grandi et al [21]. All numerical simulations were implemented in MATLAB 2017b. Ordinary differential equations were solved numerically using a variable-step, variable-order method.

### 2.2. Generation and calibration of single-cell computational models

Before evaluating the parameter–phenotype relationships, we first calibrated the atrial cell model so that majority of perturbed models would fall within the physiological range of APD at 90% of repolarization (APD_90_), maximum upstroke velocity (Vmax), and resting membrane potential (RMP). To estimate the physiological parameter range, we widely perturbed parameters using the Sobol sequence [22], that is, the 3,000 parameter combinations were log-uniformly sampled around within 25 – 400% (i.e., 2^-2^ – 2^2^) of the normal sinus rhythm (NSR) baseline which is the default setting of the Grandi model. Among 3,000 parameter combinations, 62 parameter sets showed physiological behaviors defined as APD90 of 190 – 440 ms [23], no APD alternans, RMP of −80 – −65 mV [24], and Vmax of 150 – 300 V/s [24] at steady state with pacing cycle length (PCL) of 1000 ms for 16 beats (Fig. 1A). We then defined a fine-tuned baseline as the mean parameter values within physiological conditions (Fig. 1B) and re-constructed a population of 5,000 single-cell models around within 50 – 200% (i.e., 2^-1^ – 2^1^) of the fine-tuned baseline using the Sobol sequence (Fig. 1C). Excluding non-physiological parameter combinations, a total of 603 physiological models was generated. The first 600 models were used for the downstream analysis, including ML analyses and 4-fold cross-validation (CV).

**Figure 1.**
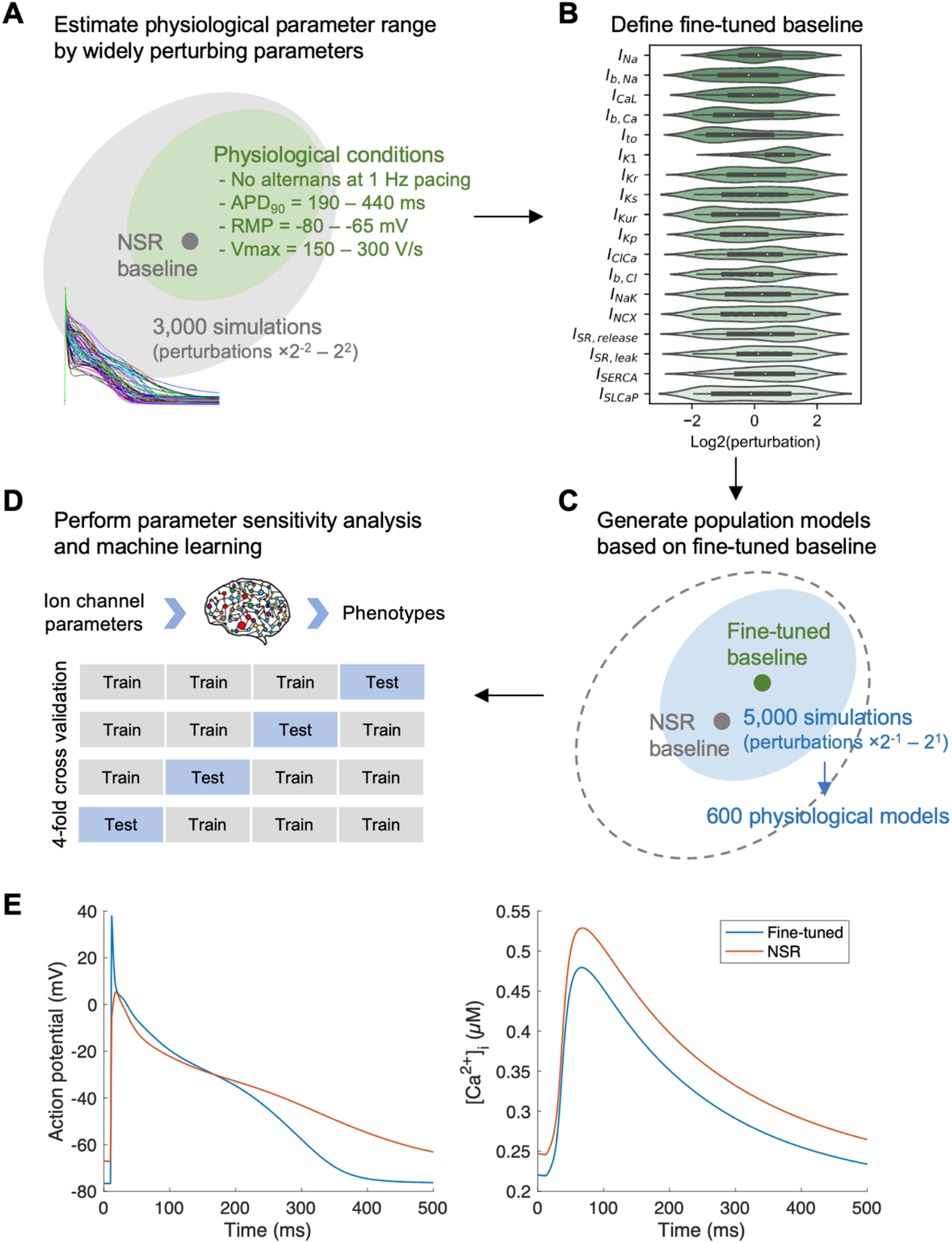
Overview of methods used in this study. (A) Using a single-cell model of human atrial cardiomyocytes, we widely perturbed ion channel parameters around the normal sinus rhythm (NSR) baseline to estimate the physiological parameter range. APD90, action potential duration at 90% of repolarization; RMP, resting membrane potential; Vmax, maximum upstroke velocity. (B) Violin plots for physiological parameter combinations. The fine-tuned baseline was defined as the mean parameter values within the physiological range (marked as white points). (C) Based on the fine-tuned baseline, we generated a population of single-cell models of atrial cardiomyocytes by log-uniformly perturbing ion channel parameters. The resultant population consists of 600 single-cell models. (D) Parameter sensitivity analysis and machine learning were used to capture the parameter–phenotype relationships at the global and local levels. Four-fold cross-validation scores were computed. (E) Action potential and intracellular calcium concentration plots of the fine-tuned baseline and NSR models at pacing cycle length of 600 ms.

### 2.3. Action potential duration restitution curve

APD restitution curves were assessed using a dynamic ramp pacing protocol: PCL of 1000, 600, 500 ms, and PCL was decreased to 300 ms in steps of 50 ms and further decreased in steps of 10 ms until failed to 1:1 capture. At each step, the cell was stimulated for 16 beats. Calcium transient amplitude (ΔCaT) was calculated as the difference between maximum and minimum values of intracellular calcium concentration. For each PCL, we evaluated APD (or calcium) alternans by determining whether the maximum and minimum APD (or ΔCaT) values in the last three beats differed by more than 2%. If APD alternans occur during pacing, we only recorded the minimum APD values in the last three beats at each PCL. The restitution curves were fitted to *APD*_90_ = *a* – *b · e^−DI/c^* where DI is a diastolic interval and *a*, *b*, and *c* are coefficients, using a nonlinear least square method [25]. The maximal slope (Smax) of the APD restitution curve was calculated as 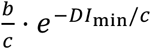 where DI_min_ is the minimum of DI.

### 2.4. Parameter sensitivity analysis

We performed a parameter sensitivity analysis to evaluate linear relationships between model parameters and various electrophysiological phenotypes, including RMP, Vmax, APD_90_, refractory period (RP), APD alternans threshold, calcium alternans threshold, and Smax. All 18 parameters were log2-transformed so that input variables follow approximately uniform distributions on (−1, 1). Since Smax shows the approximately log-normal distribution, we used log-transformed Smax values for all analyses (Fig. S1). The sensitivity of the parameter was estimated by using a multivariate linear regression method [16, 26] as *Y* = *XB* + *ϵ* where matrices *Y, X*, and *B* represent phenotypes, input parameters, and regression coefficients (i.e., sensitivity), respectively and *ϵ* is the regression error. The performance of the linear regression model was evaluated using 4-fold CV.

### 2.5. Machine learning of APD restitution curve slope

To explore the nonlinear relationships between parameters and electrophysiological phenotypes, we used random forest (RF), gradient boosting (GBM), and neural network (NN) that has one or two hidden layers. For NN models, we constructed fully-connected hidden layers with Rectified Linear Units (ReLUs) and an output layer with linear activations, and utilized a dropout regularization technique. The stochastic gradient descent algorithm was used to train the NN models. Since the performance of ML models depends critically on the hyperparameters, we used the Bayesian hyperparameter optimization algorithm with 100 iterations of optimization (see Table S1). The cost function was defined as a L2 norm of the differences between the ML-predicted values and the simulated values. All of the ML models were tested using a 4-fold CV. We then interpreted our ML model by estimating the parameter importance at both global and local levels. To assess the global parameter importance of the ML models, we computed the permutation feature importance by calculating the increase in the prediction error after randomly permuting each parameter [20]. Additionally, we evaluated parameter importance at the local level for each individual data using the local interpretable model-agnostic explanations (LIME) algorithm [19]. The individual LIME explanation values were visualized using the t-distributed stochastic neighbor embedding (tSNE) plot and clustered by the agglomerative hierarchical clustering method. All analyses were performed using the Scikit-learn, Keras, ELI5, and LIME packages of Python 3.7.

## 3. Results

### 3.1. Linear relationships for electrophysiological phenotypes

We generated the population models of 600 human atrial single-cell models by log-uniformly perturbing model parameters around the physiologically calibrated baseline (Fig. 1, see Methods for details). Briefly, we first widely perturbed the ion channel parameters around the NSR baseline to estimate the physiological parameter range and defined the fine-tune baseline as the mean parameter values within the physiological range (Fig. 1A-B). We then generated the population models around the fine-tuned baseline by log-uniformly perturbing ion channel parameters (Fig. 1C). The resultant population includes 600 single-cell models. For each single-cell model, we measured RMP, Vmax, and APD90 at 1Hz-pacing steady state, and calculated the APD restitution curve using a dynamic pacing protocol, followed by measuring APD/calcium alternans threshold and Smax. Parameter sensitivity analysis and ML models were used to explore the parameter–phenotype relationships (Fig. 1D). Steady-state action potential and intracellular calcium concentration of the baseline models are shown in Fig. 1E.

The pairwise linear correlations between electrophysiological phenotypes are shown in Fig. 2A. RMP showed a negative correlation with Vmax. Both APD_90_ at PCL of 1000 ms or 600 ms were negatively correlated with RMP and positively correlated with Vmax. There was a strong correlation between the APD alternans threshold and the calcium alternans threshold. The alternans threshold of APD_90_ or calcium showed a negative correlation with APD_90_ and a strong positive correlation with RP. We found that Smax was positively correlated with RMP, APD_90_, and RP, and negatively correlated with APD alternans threshold; however, interestingly, there was no significant correlation between Smax and calcium alternans threshold.

**Figure 2.**
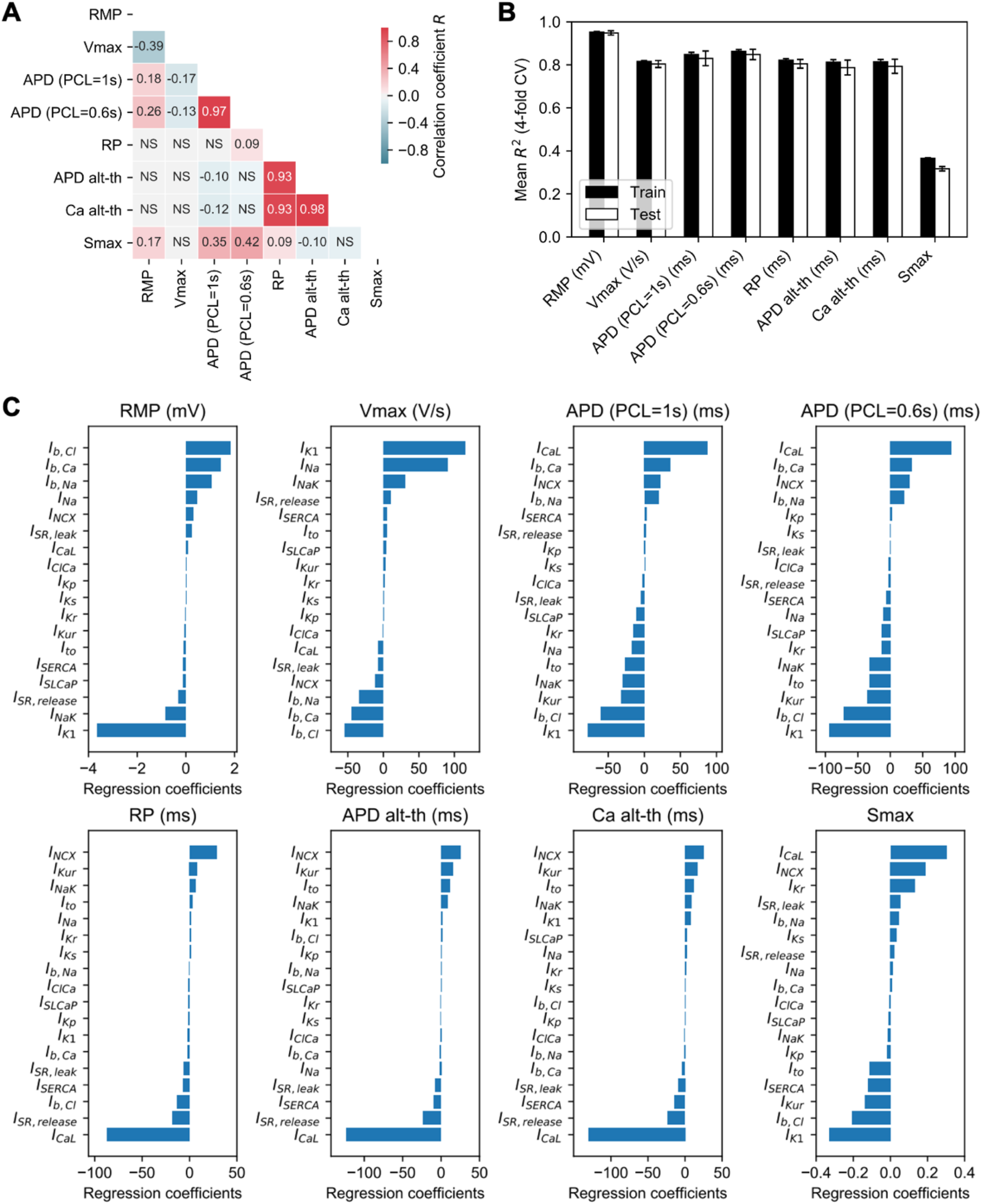
Linear relationships of electrophysiological phenotypes. (A) Pairwise correlation coefficients among electrophysiological phenotypes in 600 single-cell models. Pearson correlation coefficients were shown. NS, not significant. (B) The four-fold cross-validation scores for linear model-based parameter sensitivity analysis on electrophysiological phenotypes. Mean±SD for R^2^ values were presented. (C) Parameter sensitivities of ion channel parameters on electrophysiological phenotypes. RMP, resting membrane potential; Vmax, maximum upstroke velocity; APD, action potential duration; PCL, pacing cycle length; RP, refractory period; APD alt-th, APD alternans threshold; Ca alt-th, calcium alternans threshold; Smax, the maximal slope of APD restitution curve.

To determine how ion channel parameters affect electrophysiological phenotypes of atrial single-cell models, we performed the parameter sensitivity analysis using a linear regression model. As shown in Fig. 2B, our linear model-based sensitivity analysis could well explain the linear relationship between ion channel parameters and phenotypes except for Smax in both the training set and test set (R^2^=0.787–0.948, 4-fold CV). In contrast to the majority of phenotypes, the linear model showed poor prediction performance for Smax (R^2^=0.317±0.010, 4-fold CV). The sensitivities for electrophysiological phenotypes are shown in Fig. 2C. RMP and Vmax were primarily affected by I_K1_, I_NaK_, I_Na_, and background ion currents such as I_b,CL_, I_b,Ca_, and I_b,Na_. APD_90_ was positively correlated with I_CaL_, I_b,Ca_, I_NCX_, and I_b,Na_ and negatively correlated with I_K1_, I_b,Cl_, I_Kur_, I_to_, and I_NaK_. Both of the APD and calcium alternans thresholds depended on I_NCX_, I_Kur_, I_CaL_, I_SR,release_, I_SERCA_, and I_SR,leak_.

We found that increases in I_CaL_, I_NCX_, and I_Kr_, and decreases in I_K1_, I_b,Cl_, I_Kur_, I_SERCA_, and I_to_ are correlated with higher Smax values. To validate the results of the parameter sensitivity analysis on Smax, we perturbed each ion channel parameter predicted from the sensitivity analysis around the fine-tuned baseline and calculated APD restitution curves (Fig. 3). The *in-silico* 30%-upregulation (or downregulation) in I_CaL_, I_NCX_, and I_Kr_ resulted in steeper (or gradual) APD restitution curves, respectively (Fig. 3A-C). In contrast, *in-silico* perturbations in I_K1_, I_b,Cl_, I_Kur_, I_SERCA_, and I_to_ show inverse effects on the slope of APD restitution curves (Fig. 3D-H). These *in-silico* perturbation experiments confirmed the results of the parameter sensitivity analysis on Smax, though the linear model had poor prediction performance.

**Figure 3.**
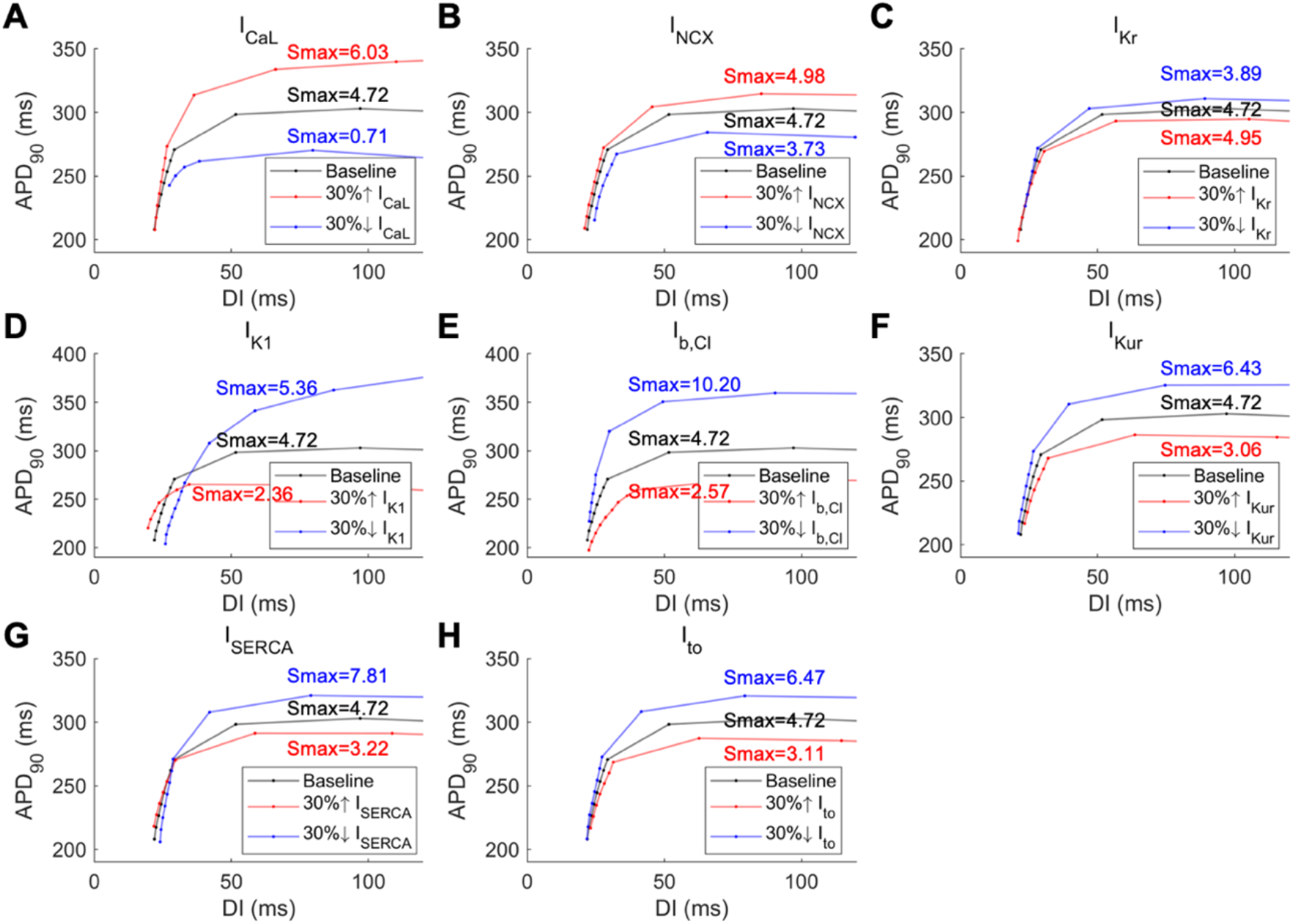
Action potential duration restitution curves after perturbing ion channel parameters. Each parameter was 30% increased or decreased from the fine-tuned baseline. Increases in I_CaL_, I_NCX_, and I_Kr_ (A-C), and decreases in I_K1_, I_b,Cl_, I_Kur_, I_SERCA_, and I_to_ (D-H) resulted in steeper APD restitution curves. APD, action potential duration; DI, diastolic interval; Smax, the maximal slope of APD restitution curve.

Furthermore, the atrial cell models were significantly clustered into four distinct groups based on the electrophysiological phenotypes (Fig. S2A). The phenotypes were significantly different between the groups (p<0.05, ANOVA test; Fig. S2B) and there were significant differences in I_Na_, I_CaL_, I_b,Ca_, I_K1_, I_b,Cl_, and I_NCX_ among the four clusters (p<0.05, ANOVA test; Fig. S2C).

### 3.2. Nonlinear analysis of APD restitution curve slope

Since conventional parameter sensitivity analysis shows poor performance to predict the parameter-Smax relationship, we trained several ML models better to explain the nonlinear relationships between parameters and Smax. For each ML model, we optimized hyperparameters using the Bayesian hyperparameter optimization algorithm (Table S1) and calculated 4-fold CV scores to evaluate the prediction performance (Fig. 4A). Two ensemble learning models show moderate prediction performances, which are slightly higher than the linear model (R^2^=0.388±0.025 for RF and 0.417±0.025 for GBM, 4-fold CV). The neural network models had better prediction performance than ensemble learning models (R^2^=0.495±0.034 for the one-hidden-layer model and 0.502±0.023 for the two-hidden-layer model, 4-fold CV). Among our ML models, the NN with two hidden layers showed the best prediction performance (Fig. 4B); thus, we chose this NN model for the downstream analysis.

**Figure 4.**
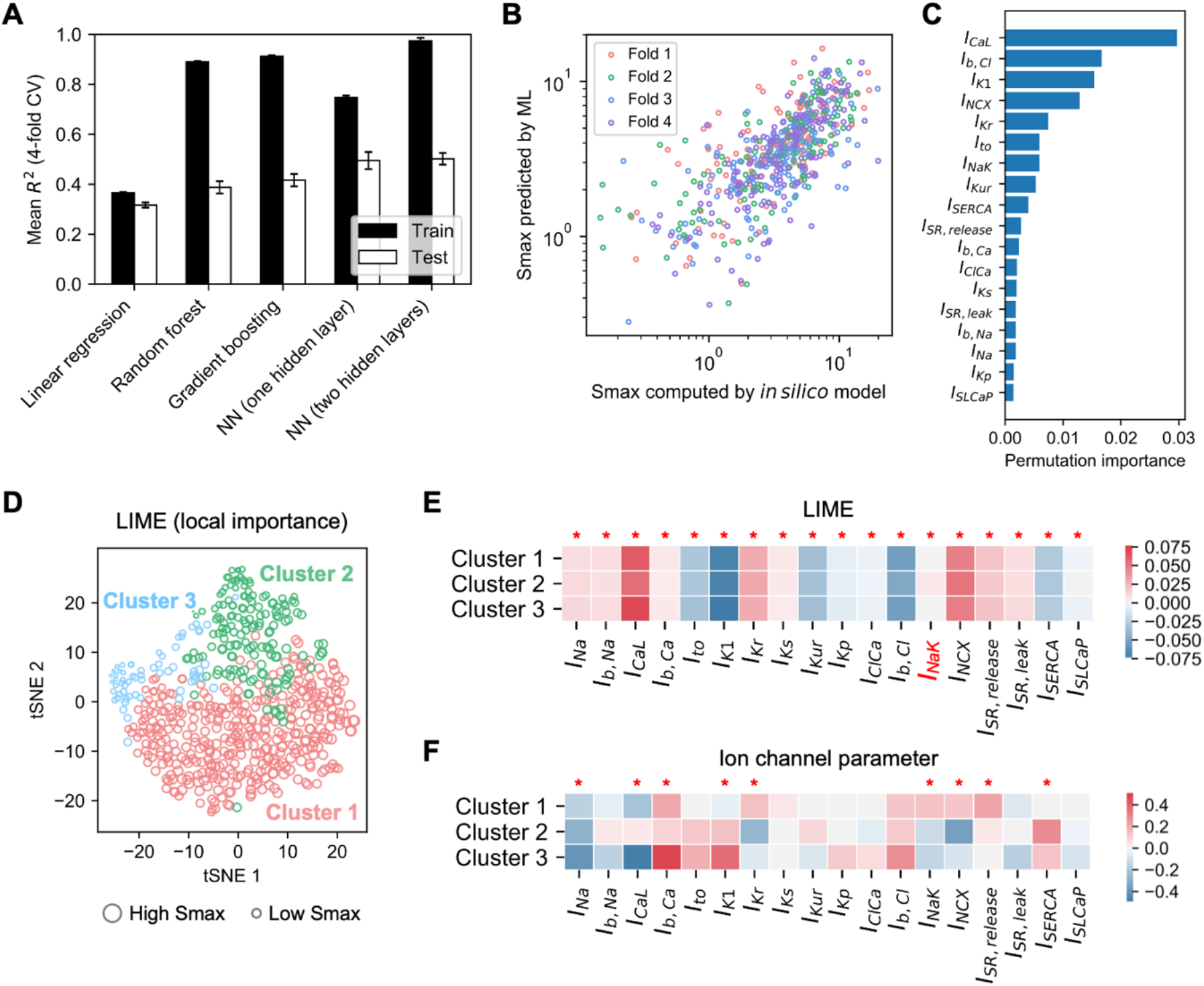
Machine learning of the maximal slope of the APD restitution curve (Smax). (A) Linear regression, random forest, gradient boosting, and neural network (NN) models were used to train parameter–Smax relationships. Mean±SD for 4-fold cross-validation scores (R^2^) were presented. (B) The correlations between Smax values simulated by *in silico* models vs. those predicted by the NN model with two hidden layers. (C) Global parameter sensitivity was computed as permutation feature importance. (D) Local parameter importance was computed for each data point using the LIME method and visualized using a tSNE plot. Data were clustered into three groups based on the local importance profiles. The size of the data point indicates Smax values. (E) Mean LIME values were visualized for each cluster. At each parameter, LIME values were significantly different across the clusters. (F) Mean ion channel parameters were visualized for each cluster. The parameters that were significantly different across the clusters were marked. *p<0.05.

To interpret the ML model, we evaluated the parameter importance at the global and local levels by computing the permutation feature importance and the LIME explanation values, respectively. As shown in Fig. 4C, I_CaL_, I_b,Cl_, I_K1_, I_NCX_, I_Kr_, I_to_, I_NaK_, and I_Kur_ were assigned high global permutation importance values. One interesting finding is that I_NaK_ plays a significant role in determining Smax as well as I_to_ and I_Kur_ (Fig. 4C), while the linear model could not capture the association between I_NaK_ and Smax (Fig. 2C). Additionally, we visualized the individual local parameter importance (i.e., LIME) using the tSNE method. As it can be seen from Fig. 4D, data points are correlated well with Smax values, indicating that the ML model is able to provide a reasonable prediction of Smax. Moreover, population data could be clustered into three distinct groups based on the hierarchical clustering of LIME profiles. For each feature, the LIME values were significantly different among the three clusters (p<0.05, ANOVA test; Fig. 4E). While most parameters showed similar directions/magnitude of the local importance, I_NaK_ had negative LIME values in the clusters 1/3, but positive values in the cluster 2. We further compared the log2-values of ion channel parameters among the three groups. There were significant differences in I_Na_, I_CaL_, I_b,Ca_, I_K1_, I_Kr_, I_NaK_, I_NCX_, I_SR,release_, and I_SERCA_ between the clusters (p<0.05, ANOVA test; Fig. 4F).

### 3.3. Role of Na-K pump in APD restitution curve

The APD_90_ at PCL of 600 ms and APD/calcium alternans thresholds were significantly different across the three LIME-based clusters (p<0.05, ANOVA test). Steady-state action potential and CaT of the average models of the clusters are shown in Fig. 5A, indicating the alterations in the baseline APD and CaT.

**Figure 5.**
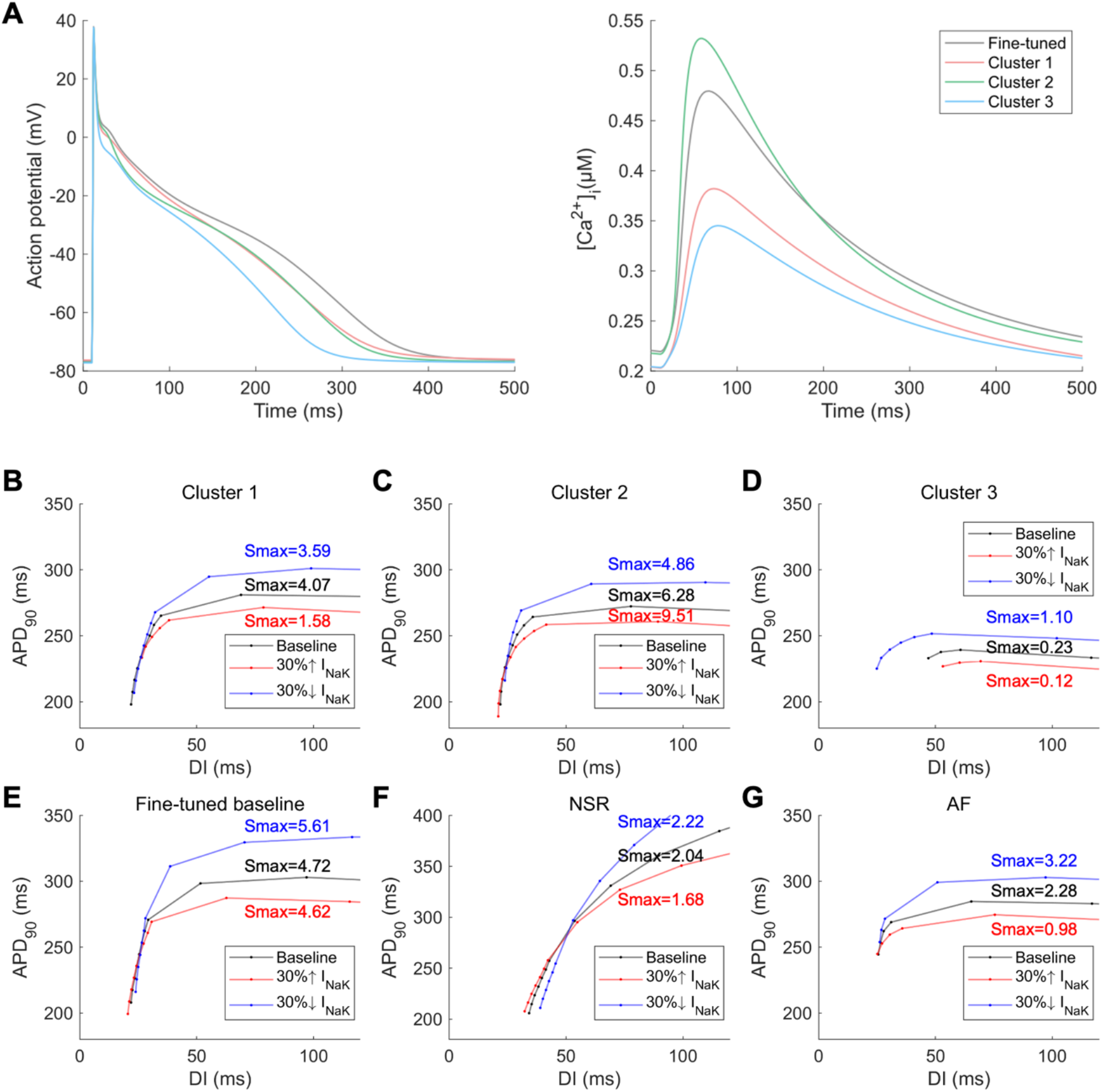
(A) Action potential and calcium transient plots of the fine-tuned baseline model and the average models for the LIME-based clusters. Steady-state action potentials are shown at pacing cycle length of 600 ms. (B-G) Action potential duration restitution curves were calculated using a dynamical pacing protocol after perturbing Na-K currents. The Na-K currents were 30% increased or decreased from the average parameter set for each LIME-based cluster (B-D), and the fine-tuned baseline (E), normal sinus rhythm (NSR) condition (F), and atrial fibrillation (AF) condition (G). APD, action potential duration; DI, diastolic interval; Smax, the maximal slope of APD restitution curve.

To validate the findings from the ML model, we performed *in silico* experiments in the single-cell model by perturbing I_NaK_ around the mean parameter set of each cluster (Fig. 5B-G). In the cluster 1, both 30%-upregulation and downregulation in I_NaK_ decreased Smax (Fig. 5B). However, the clusters 2 and 3 showed concordant results with the predictions from the LIME analysis. In the cluster 2, the increase (or decrease) in I_NaK_ resulted in the higher (or lower) Smax, respectively (Fig. 5C). In contrast to the cluster 2, the perturbations in I_NaK_ showed inverse effects on Smax in the cluster 3 (Fig. 5D). Additionally, we perturbed the Na-K current around the fine-tuned baseline, NSR, and AF baselines. The AF baseline was defined as Grandi et al [21], which decreased I_CaL_, I_to_, I_Kur_ and I_SERCA_, and increased I_K1_, I_Ks_ and I_NCX_. For all the fine-tuned baseline, NSR, and AF baselines, the *in-silico* 30%-upregulation (or downregulation) in I_NaK_ resulted in the decreases (or increases) in Smax, respectively (Fig. 5E-G).

## 4. Discussion

By performing linear model-based parameter sensitivity analyses on the single-cell atrial computational models, we determined the effects of individual ion channels on the electrophysiological phenotypes, including RMP, Vmax, APD_90_, RP, APD alternans threshold, calcium alternans threshold, and Smax. In addition, we developed the ML models for predicting Smax from individual ion channel profiles, which has better prediction performance than the linear model. We discovered the biphasic behavior of I_NaK_ in determining Smax, which was not appreciated by conventional linear model-based parameter sensitivity analyses.

Smax is one of the key factors to determine single cell-level dynamic instability of cardiomyocytes and is closely linked with rate-dependent intracellular Na^+^ and Ca^2+^ accumulations, which are directly or indirectly controlled by I_NaK_, I_NCX_, I_Ks_, I_to_, I_CaL_, and SR currents [27–29]. However, how various electrical remodeling conditions affect the APD restitution kinetics has been poorly understood. Our systems biology approaches successfully identified various subpopulations of atrial cell models that show heterogeneous APD restitution kinetics in response to the perturbations in I_NaK_. The subcellular dynamical mechanisms underlying heterogeneous APD restitution curve behaviors should be further investigated.

Conventional parameter sensitivity analysis could determine the linear relationships between ion channel profiles and ‘static’ arrhythmia markers, such as APD, RMP, Vmax, and alternans threshold [30]. Recently, several ML approaches have been utilized to evaluate the arrhythmogenicity from electrical/structural remodeling status [31–34]. We previously demonstrated that statistical physics-derived methods, such as a maximum entropy model, can intuitively describe functional coherence properties in the spatiotemporal wave dynamics of cardiac fibrillation [35]; however, those methods typically require huge computational costs for systemic population-scale data analyses. In this paper, our interpretable ML approaches demonstrated the nonlinear relationships between ion channel parameters and ‘dynamic’ arrhythmia markers, especially Smax, at both local and global levels on the parameter space. Although we only explored single-cell cardiac models in this paper, our ML approaches have potential generalizability to determine the nonlinear relationships between model parameters and heterogeneous dynamic phenotypes and drug response patterns in multiscale mathematical models [36, 37]. Additionally, deep learning models might be helpful to better understand parameter–phenotype mapping in cardiac systems [38]. We also expect that our combined top-down and bottom-up computational approaches can be extended to provide functional (physiological) interpretations of single-cell omics dataset [39, 40] by integrating gene/protein expression and interaction information at the single-cell resolution [41].

The uncertainty and variability in mathematical models of cardiac electrophysiology are inevitable and may lead to inaccurate outcomes in computational modeling-assisted clinical procedures [42, 43]. The multiscale and high-dimensional nature of heart models, including single-cell atrial models used in our study, further amplify such model uncertainty and variability. Our interpretable and explainable ML approaches could systemically describe nonlinear parameter–phenotype relationships in high-dimensional spaces. Our unbiased systems biology approaches have potential generalizability to quantify the parameter uncertainty in mechanism-based dynamical models. Probabilistic models, such as restitution curve emulators, might be helpful in identifying the model uncertainty and calibrating parameter combinations in *in silico* heart models [44].

The APD restitution curves and Smax are affected by pacing protocol-dependent memory effects and cannot fully reflect dynamic instability in tissue-level phenomenon [45–47]. These computational and physiological uncertainties of Smax might contribute to the poor prediction performance of Smax in our ML models. Other robust various electrophysiological parameters as well as structural parameters, such as fibrosis [48, 49], tissue thickness [50], and curvature [51], should be further investigated for systemic evaluations of pro- or anti-arrhythmogenic conditions in both single-cell and tissue-level models. To further validate our ML models, external validation using experimental/clinical data or other mathematical models should be considered.

## Abbreviations and Acronyms

APD: action potential duration
CaT: calcium transient
DI: diastolic interval
I_b,~_: background current
I_CaL_: L-type calcium current
I_K1_: inward rectifier potassium current
I_Kr_: rapidly activating delayed rectifier potassium current
I_Ks_: slowly activating delayed rectifier potassium current
I_Kur_: ultrarapid delayed rectifier potassium current
I_Na_: fast sodium current
I_NaK_: sodium-potassium pump current
I_NCK_: sodium-calcium exchange current
I_SERCA_: sarcoplasmic reticulum calcium pump current
I_SLCaP_: sarcolemma calcium pump transport
I_to_: transient outward potassium current
LIME: local interpretable model-agnostic explanations
ML: machine learning
NSR: normal sinus rhythm
PCL: pacing cycle length
RMP: resting membrane potential
RP: refractory period
Smax: the maximal slope of APD restitution curve
SR: sarcoplasmic reticulum
tSNE: t-distributed stochastic neighbor embedding
Vmax: maximum upstroke velocity

## Acknowledgments

This research was presented at the *66^th^ Biophysical Society Annual Meeting*, San Francisco, California, 2022. This work was supported by the National Research Foundation research grant funded by the Ministry of Science and ICT (2021R1F1A1063218 to Y.-S. Lee). We would like to thank K. Park for insightful discussions on the dynamic systems biology modeling.

## Authorship Contribution

**E. Song:** Conceptualization, Methodology, Software, Investigation, Data curation, Visualization, Writing – original draft, Writing – review & editing. **Y.-S. Lee:** Supervision, Writing – review & editing, Funding acquisition.

---

**Table S1.** The optimal hyperparameters of machine learning models based on the Bayesian hyperparameter optimization method with gaussian processes.

| Model | Hyperparameter | Range | Optimal value |
| --- | --- | --- | --- |
| RF | Max depth | [3, 100] | 44 |
|  | Min samples split | [2, 100] | 2 |
|  | Min samples leaf | [1, 50] | 3 |
|  | Number of estimators | [10, 1000] | 758 |
| GBM | Max depth | [3, 15] | 10 |
|  | Min samples split | [2, 100] | 98 |
|  | Min samples leaf | [1, 50] | 46 |
|  | Number of estimators | [10, 1000] | 919 |
|  | Learning rate | [0.001, 1.0] | 0.0270 |
| NN<br>(one hidden layer) | Size of hidden layer | [1, 100] | 67 |
|  | Learning rate | [0.001, 0.5] | 0.0786 |
|  | Batch size | [5, 18] | 18 |
|  | Epochs | [50, 500] | 466 |
| NN<br>(two hidden layers) | Size of hidden layer 1 | [1, 100] | 99 |
|  | Size of hidden layer 2 | [1, 100] | 34 |
|  | Dropout rate | [0.0, 0.5] | 0.1336 |
|  | Learning rate | [0.001, 0.5] | 0.2482 |
|  | Batch size | [5, 18] | 8 |
|  | Epochs | [50, 500] | 499 |

RF, random forest; GBM, gradient boosting; NN, neural network.

**Figure S1.**
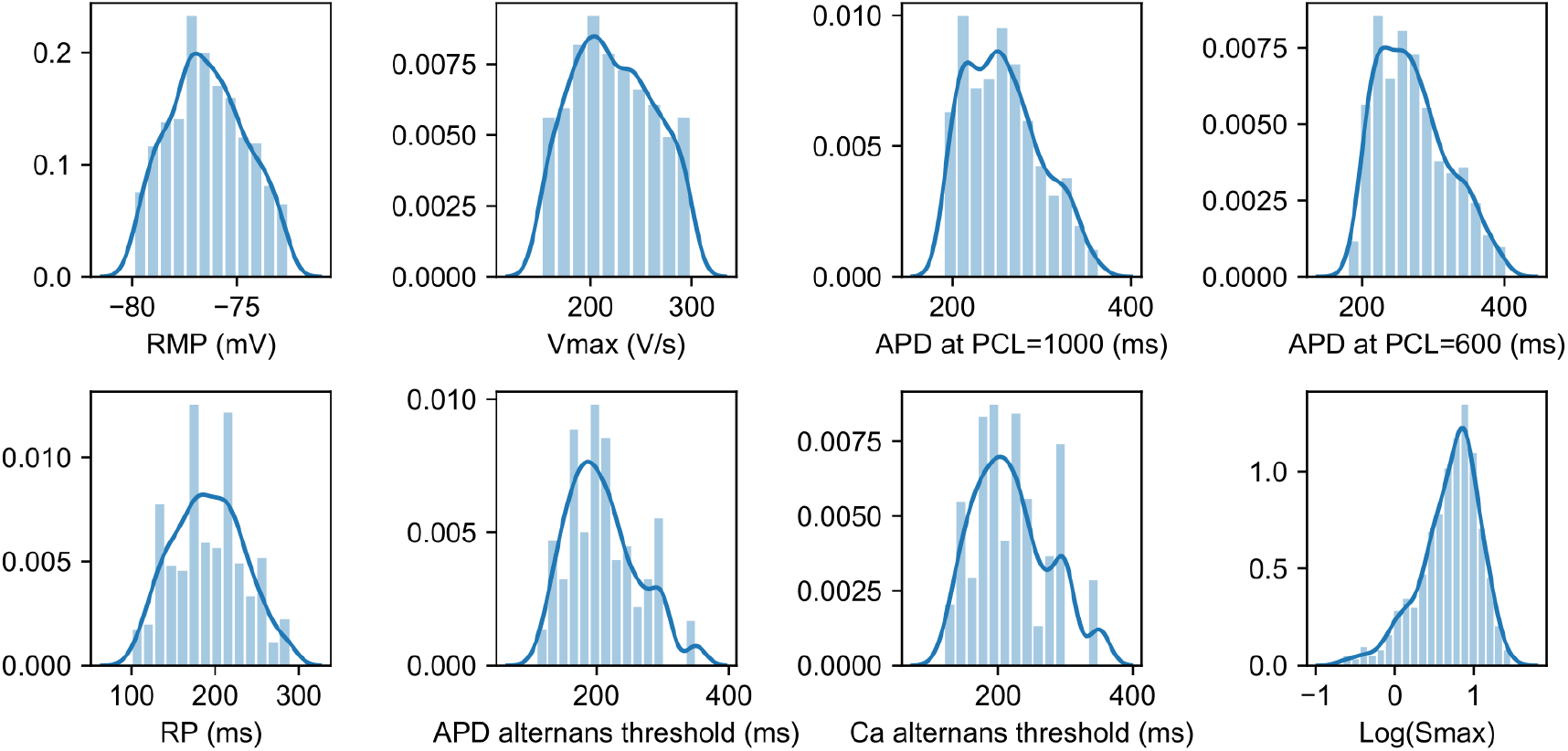
Distributions of electrophysiological phenotypes of the atrial cell models.

**Figure S2.**
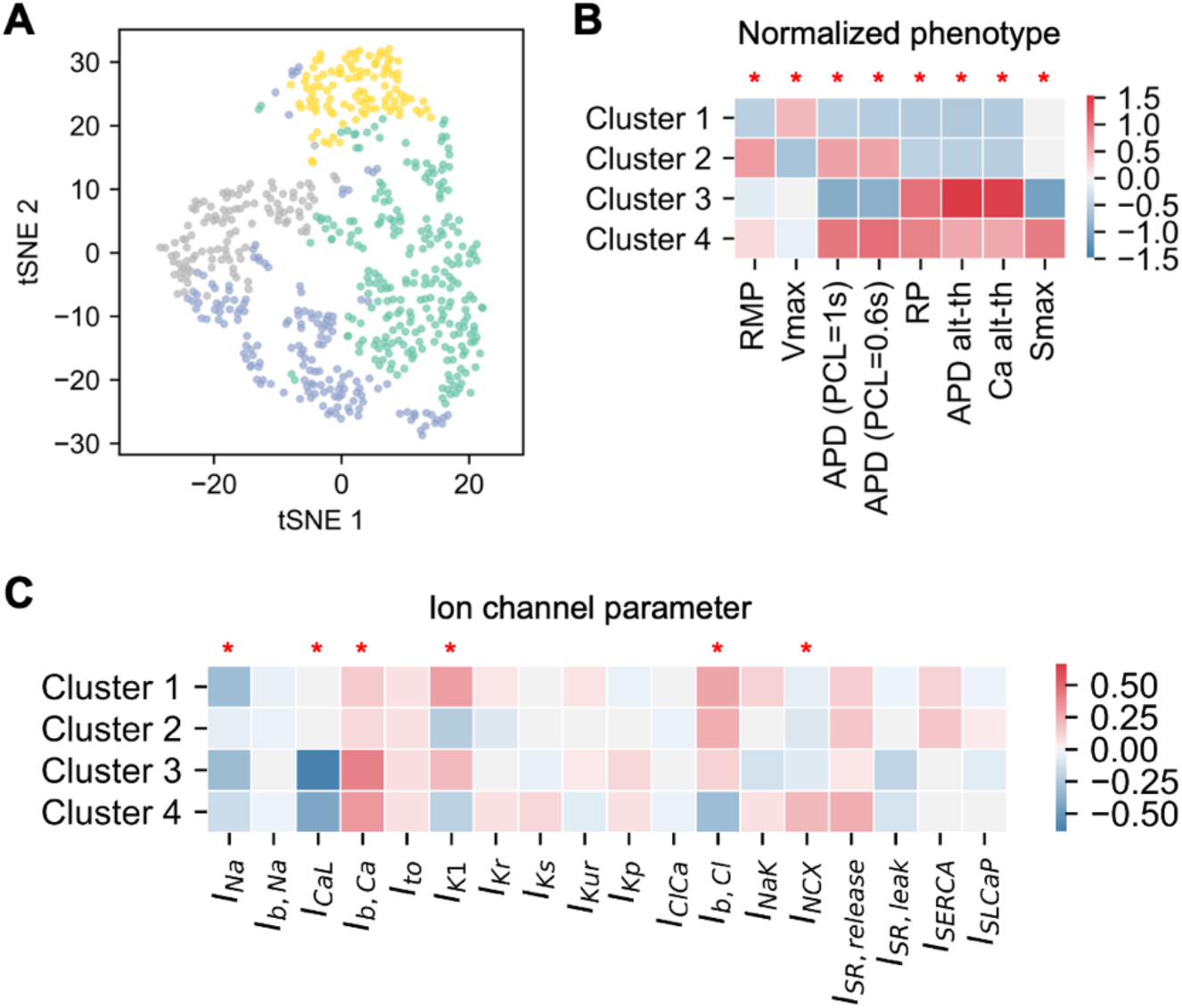
(A) Electrophysiological phenotype sets were visualized and clustered into four groups using a tSNE plot. (B) Mean phenotype values were visualized for each cluster. (C) Mean ion channel parameter values were visualized for each cluster. The parameters that were significantly different across the clusters were marked. *p<0.05.

